# Deciphering dermal fibroblast behavior in 3D bioprinted dermis contructs

**DOI:** 10.1101/2023.03.07.531460

**Authors:** Laura Chastagnier, Naima el-Kholti, Lucie Essayan, Céline Thomann, Edwin-Joffrey Courtial, Christophe Marquette, Emma Petiot

## Abstract

In recent years, numerous strategies have emerged to answer the growing demand for graftable tissues. Tissue engineering and *in-vitro* production are one of them. Among all the engineered tissues, skin is one of the most advanced. Nevertheless, biofabrication of graftable and fully functional skin substitutes is still far from being reached. Skin reconstruction, particularly dermis, necessitates cultivation and maturation for several weeks (> 3 weeks) to recover the tissue’s composition and functions, which prevent its transfer to clinical applications. Thus, several strategies, including 3D bioprinting, have been explored to accelerate these productions. In the present study, based on the successful application of 3D bioprinting achieved by our group for skin reconstruction in 21 days, we propose to detail the biological behaviors and maturation phases occurring in the bioprinted skin construct thanks to a descriptive approach transferred from the bioprocess field. The aim is to comprehensively characterize dermis construct maturation phases (cell proliferation and ECM secretion) to master later the interdependent and consecutive mechanisms involved in *in-vitro* production. Thus, standardized quantitative techniques were deployed to describe 3D bioprinted dermis proliferation and maturation phases. Then, in a second step, various parameters potentially impacting the dermis reconstruction phases were evaluated to challenge our methodology and reveal the biological behavior described (fibroblast proliferation and migration, cell death, ECM remodeling with MMP secretion). The parameters studied concern the bioprinting practice including various printed geometries, bioink formulations and cellular physiology in relation with their nutritional supplementation with selected medium additives.

## Introduction

Tissue engineering applied to regenerative medicine aims to develop implantable tissues to overcome the recurrent graft shortage and rejection issues. Numerous tissues were cultivated *in vitro* – skin[1],cornea [2], retina [3], bone [4] cartilage [5]myocardium [6], liver [7]or musculoskeletal tissues [8] – but, the only validated clinical application is the autologous epidermis to treat severe deep burnt patients [9]. Up to now, all the other tissues, including the full-thickness skin models, were applied only for *in vitro* testing purposes, such as cosmetology and toxicology assays [10]. Regarding clinical applications, grafting capabilities depend strongly on the tissue’s capacity to enable stitching and anastomosis. Reaching appropriated mechanical properties is thus crucial and still represents the bottleneck on which the tissue engineering community is striking. In this quest to produce implantable tissue, skin is considered the most advanced application.

Currently, using a 3D scaffold-based approach such as “collagen sponges”, the production of a differentiated and mature dermis lasts approximately 28 days [11]. This is a substantial duration to which must be added the necessary fibroblast pre-culture after extraction from a biopsy. Additionally, in the case of skin production, the epidermalization step of the mature dermis brings the total tissue production to 45 to 56 days [11].

The recent use of 3D bioprinting significantly reduced this time to obtain dermis substitutes. Thus, it is now possible to reach full-thickness skin within 21 days using extrusion-based 3D bioprinting [12]. In this method, 3-dimensional complex tissue structures are produced by the successive deposition of layers of cellularized proliferative bioink, leading to homogeneous cell distribution [1]. Previous works of our group demonstrated that over these 21 days maturation period, the proliferative bioink is ultimately degraded and replaced by the fibroblast neo-synthesized extracellular matrix (ECM) where type I and V, collagen, vimentin, fibrillin, and elastin are abundantly expressed [1,12]. Indeed, it is well known that during dermis production, a maturation step is required to enable extracellular matrix (ECM) secretion. This matrix is composed of a complex network of macromolecules (proteins, glycosaminoglycans, proteoglycans, glycoproteins) [13] where collagen, one of the fibrous proteins secreted by fibroblasts (collagens, elastin, fibronectin and laminin), represents 70% of the dermis dry mass [14].

The biological behaviors involved in a bioprinted dermis tissue maturation process is close to the *in-vivo* wound healing process which is split in a fibroblast migration and proliferation phase (i) prior to their ECM secretion (ii) and a subsequent tissue contraction phase (iii) (**Figure 1**). This ECM remodeling is a central process in wound healing and scarring. It consists of the ECM components degradation to favor cell proliferation and migration followed by neo-synthesis of new matrix components. Several enzymes are involved in these processes, and here, vitamin C is a mandatory cofactor of prolyl and lysyl-hydroxylases. These two enzymes are involved in the synthesis of collagen through hydroxylating the proline and lysine residues of the peptides forming the collagen filament (α-chains), thus allowing the formation of hydrogen bonds between the α-chains [15]. Vitamin C addition during *in vitro* skin maturation is thus the gold standard medium supplement to trigger ECM synthesis [16].

**Figure 1.**
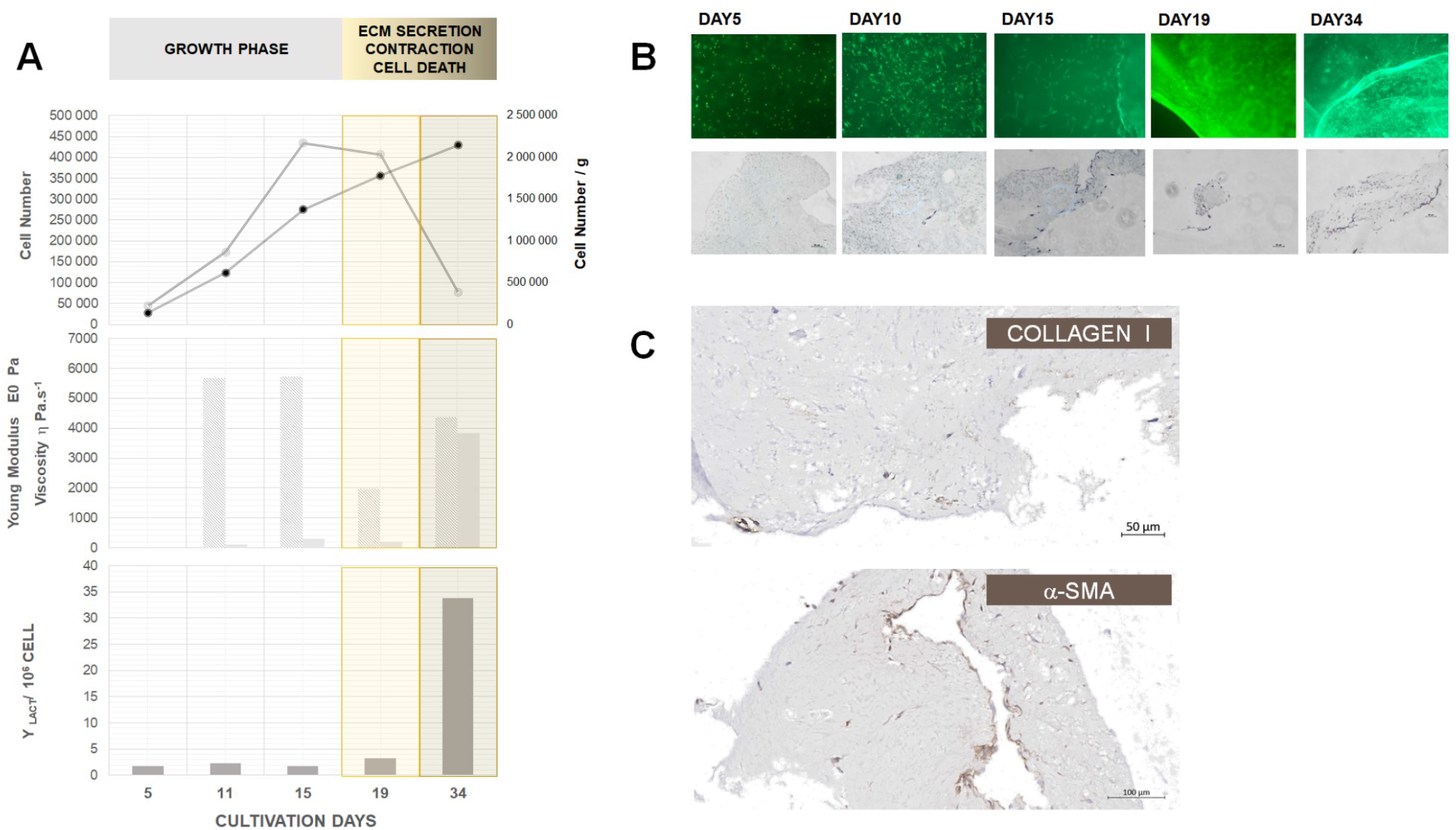
A- Proliferation profile (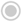 total cell number in the construct; ● cell number per grams of constructs) associated with mechanical characterization (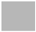 Viscosity 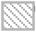, Young modulus) and metabolic activity (lactic acid secretion) of the 3D bioprinted constructs; B-Bioprinted construct cellularization evolution over time: upper part, monitoring of viable cells by calcein-green fluorescent staining, lower part, histological analysis with Masson trichrome staining. Cells are identified in pink. C- Immuno-histological staining of Collagen I secretion and a-SMA proteins at day 15 post-printing. The bioink formulation used for this study comprises 5% Gelatin, 2% Alginate and 2% fibrinogen (Formulation A).

On the opposite, several biological mechanisms allow the homeostasis of the ECM by activating or inhibiting enzymes responsible for the degradation of ECM compounds: the matrix metalloproteinases (MMPs) [17–19]. These enzymes are secreted by fibroblasts or myofibroblasts (i.e. differentiated contractile fibroblasts) [20]. In healthy tissue, the metalloproteases are produced as inactive form (a pro-domain blocks access to the catalytic site) [21]. However, when secreted in large quantities, these enzymes can cleave the pro-domains, leading to increased degradation of the extracellular matrix. Thus, they play an important role in ECM remodeling and cell migration during wound healing and scaring [21,22]. *In vitro* and *in vivo*, the activity of these enzymes has already been controlled by adding Tissue Inhibitor of Metalloproteinases (TIMP), such as the pharmacological inhibitors Batimastat® and Prinomastat® [23,24]. In the present study, we aimed at reaching an in-depth characterization of dermis constructs maturation phases (cell proliferation and ECM secretion). Indeed, reconstituting functional dermis and particularly its mechanical properties and fibro-elastic characteristics is not trivial. Thus, it is of high importance to understand and master the interdependent and consecutive mechanisms involved in wound healing to produce *in-vitro* dermis. The applied strategy is inspired from bioprocess monitoring and characterization strategies to reach biological mechanism comprehension. The first part of the study consists in developing standardized quantitative techniques to describe 3D bioprinted dermis proliferation and maturation phases. The second part focus on evaluating various parameters potentially impacting the dermis reconstruction phases to validate our methodologies. Parameters evaluated were the printed geometries, the bioink formulation and the supplementation of our cultivation medium with selected additives. Two growth factors (FGF & EGF) known for their action on fibroblasts’ proliferation and migration [25,26] were evaluated as well as two pharmaceutical MMP inhibitors (Prinomastat® and Batimastat®).

## Materials and methods

### Cell origin and cultivation

Human fibroblasts extracted from dermis biopsy were graciously provided by Labskin Creations (Lyon, France). The human dermal fibroblasts were cultured in a reconstituted Complete medium composed of DMEM medium (Dulbecco’s Modified Eagle Medium Gibco, Thermo Fisher Scientific, USA, 31966-021) supplemented with 10% (v/v) fetal calf serum (FCS, Gibco Cell Culture, 10270-106), 0.5% (v/v) Penicillin-Streptomycin antibiotics at 10,000 U/mL (Gibco, Thermo Fisher Scientific, 15140122), and 0.4% (v/v) antifungal (Amphotericin B, Gibco, Thermo Fisher Scientific, 15290-018). Cell cultures were incubated at 37°C in 5% CO_2_ incubators. The cells were grown in T-flasks and sub-cultured once a week when reaching confluency. Cell culture supernatants at confluency, after 7 days of culture were collected and used as conditioned media.

### 3D Bioprinting and dermis construct cultivation

The studied human dermis was produced through 3D bioprinting of a bioink cellularized with fibroblasts. The already published and patented bioink [1] consists of a combination of porcine gelatin (Gelatin from porcine skin, Sigma Aldrich, United States, G1890), alginate (Alginic acid salt very low viscosity, Alfa Aesar, United States, A18565) and fibrinogen (Fibrinogen from bovine plasma, Sigma Aldrich, F8630). Two formulations of the bioink were tested. Different ratios between gelatin and alginate allowed to reach different mechanical and bioprintability properties of the biomaterial. Thus, Bioink formulation A consisted of 5% (w/v) Gelatin, 2% (w/v) Alginate and 2% (w/v) fibrinogen while formulation B consisted of 10% (w/v) of gelatin, 1% (w/v) of Alginate and 2% (w/v) of fibrinogen. Bioinks were seeded with 2D cultured fibroblasts at a concentration of 0.125×10^6^ cells/ml. The 3D bioprinting was performed using the 6-axis robotic BioassemblyBot® bioprinter (Advanced Solutions Life Sciences, USA) equipped with a controlled atmosphere enclosure (HEPA filter). According to the bioinks rheological behaviors and to use similar flow rates, formulation A and B were printed at 21°C and 28°C, respectively. Standard operational parameters were a pneumatic extrusion tool loaded with a 10CC cartridge (Nordson EFD), mounted with a 400 µm diameter (6.3 mm long) nozzle (Nordson EFD), and an applied extrusion pressure of 20 PSI. Once bioprinted, the dermis construct was consolidated in a solution composed of 3% (w/v) calcium chloride (Calcium Chloride, Sigma Aldrich, C5670) and Thrombin at a concentration of 10 U/mL (Thrombin from bovine plasma, Sigma Aldrich, T4648-10KU). Tissue constructs were bioprinted with different porosity and thickness as presented in **Table 1**. A slab geometry was chosen for these experiments. The design was realized thanks to the TSIM interface (Advanced Solutions Life Sciences, USA).

Standard cultivation protocol of 3D bioprinted dermis constructs was performed in 6-well plates at 37°C in 5% CO_2_ incubator within 3 ml of complete medium. Media supplements tested were FGF-2 at a concentration of 10 ng/ml (Human FGF-2 IS premium grade, MACS Miltenyi Biotec, Germany, 130-104-922), EGF at a concentration of 10 ng/ml (Recombinant Human EGF, Gibco, Thermo Fisher Scientific, PHG0314), Prinomastat® at a concentration of 10 nM [24] (Sigma Aldrich, PZ0198), Batimastat® at a concentration of 1 μM [27] (Sigma Aldrich, SML0041) and vitamin C at a concentration of 50 μg/mL [16] (L-ascorbic acid 2-phosphate sesquimagnesium salt hydrate, Sigma Aldrich, A8960). For the study of the effect of the tissue 3D geometry, culture medium was renewed twice a week.

### Bioprinted dermis characterization

#### Cell proliferation

The cell network was observed thanks to calcein green staining. Bioprinted constructs were incubated for 30 minutes at 37°C in the presence of 1 μM Calcein AM (Invitrogen, Thermo Fisher Scientific, C1430) before being observed under a fluorescence microscope (Nikon Eclipse Ts2R). Cell proliferation trends were described thanks to cell counting after bioprinted construct dissociation. Cells dissociation protocol was carried out by incubation for 5 minutes at 37°C in a solution of trypsin-EDTA 0.05% in PBS (Gibco, Thermo Fischer Scientific, 14190-094) followed by 10 minutes at 37°C in a 3% (w/v) collagenase A solution in PBS (Collagenase A, Roche Diagnostics GmbH, Germany, 10103586001). Cells were recovered by centrifugation for 5 minutes at 300g prior cell counting on Malassez cells after Trypan blue staining. Duplicated samples allowed to evaluate a mean standard error of 6% on cell counts.

The maximal growth rate in 3D constructs (µ_3D_) was determined during the exponential growth phase by plotting ln(X)=f(t) where X is the cell number. The value of µ_3D,_ expressed in day^-1^, correspond to the slope of the curves. Then, the population doubling time (PDT) was calculated thanks to the following equation:

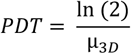

The metabolic activity was evaluated by quantification of the L-lactic acid concentration in the culture supernatant with a commercial assay kit (L-Lactic Acid (L-Lactate) Assay Kit, K-LATE, Megazyme) according to manufacturer’s instructions. The L-lactic acid production yield (Y_lact/X_) was calculated with the following formula where C_lact_ is the lactate concentration in g/l and X is the cell concentration:

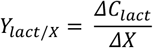

Error bars were calculated as the sum of respective errors from lactate quantification and cell numeration as follow :

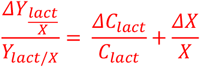

### Bioprinted construct contraction

Contraction of the bioprinted dermis was followed by recording of macroscopic evolution and expressed as shrinkage factor (%) already described in Pragnere et al. (2022)[28] It was evaluated thanks to image analysis using ImageJ software (NIH, Version 1.46). The total area of the construct was measured from a macroscopic construct picture, standardized to the area of the culture well. The ratio (area of the tissue/well area) obtained was then compared to the one measured just after bioprinting and consolidation (at day 1) to determine the following shrinkage factor, with measurement error of 0.9 %:

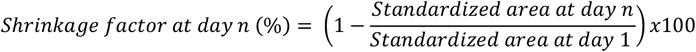

### Histological staining

For histological staining, constructs were fixed using paraformaldehyde during one night at 4°C (AntigenFix, Diapath). They were then dehydrated in a gradient of ethanol and embedded in paraffin. Samples were cut into 5 µm thick sections. For Masson’s Trichrome staining, sections were deparaffinized but not rehydrated. Staining was performed using GROAT’s hematoxylin for 30 seconds followed by Ponceau-Fuchsin solution (0.07% (w/v) Ponceau Xylidine, 0.03% (w/v) Fuchsin acid, 0.2% (v/v) glacial acetic acid) phosphomolybdic acid-orange G (4% (w/v) phosphomolybdic acid, 2% (w/v) orange G) and light green (0.2% (w/v) light green, 0.2% (v/v) glacial acetic acid) incubation for 3 minutes. Protocol for immuno histochemical staining of both α−SMA and Collagen I proteins were previously described in [28]Sections were finally dehydrated and mounted using Bioumount DPX (Biognost BM500).

### Electron Microscopy

Dermis constructs were cut into small pieces and fixed in 2% glutaraldehyde for 2 hours at 4°C. Samples were washed three times for 1 hour at 4°C and postfixed with 2% OsO_4_ for 1hour at 4°C. Then, tissues were dehydrated in graded series of ethanol and transferred to propylene oxide. Impregnation was performed with Epon A (75%) plus Epon B (25%) plus DMP30 (1.7%). Inclusion was obtained by polymerization at 60°C for 72 hours. Sections were made on a UC7 (Leica) ultramicrotome. Area’s alterations were first selected on semi-thin sections (1000 nm thick) stained with methylene blue azur II. Ultrathin sections (approximately 70 nm thick) were cut, mounted on 200 mesh copper grids coated with 1:1000 poly-lysine, stabilized for 1 day at room temperature (RT) and contrasted with uranyl acetate and lead citrate. Sections were examined with a Jeol 1400JEM (Tokyo, Japan) transmission electron microscope, 80Kv, equipped with Orius 600 and Orius 1000 cameras and digital micrograph.

#### DMA measurements

The viscoelastic behavior of bioink was characterized by frequency sweep experiments in dynamic mechanical analysis (DMA) in compression mode. These experiments were conducted with a rotational rheometer (DHR2, TA Instruments, Guyancourt, France) with a DMA mode (torque = 0N) using disk-shaped samples and a parallel plate geometry (8 mm). A preliminary study was performed to define the linear viscoelastic domain, which corresponds to the displacement range where the material properties are assumed to be constant. This domain is determined using oscillatory compression experiments with constant frequency and varying displacement. Then, dynamic compression tests were performed with a frequency range of 0.1 to 10Hz (*i*.*e*. 0.628 to 62.8 rad/s) at a constant displacement, which is within the linear viscoelastic regime. In these dynamic compression tests, bioink undergoes a periodical mechanical strain *ε* of very small amplitude *ε*_0_ (< 1%) and of angular frequency *ω* following the equation (1):

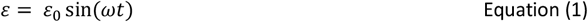

In the case of the generalized Maxwell model, the storage E′(ω) part of the complex modulus is expressed by the equations (2):

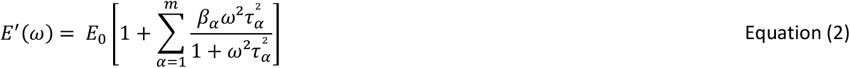

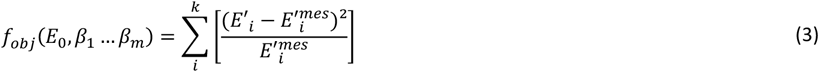

where *E*_0_ is Young’s modulus of the isolated spring. The relaxation times, *τ*_*α*_, and the dimensionless reference parameters *β*_*α*_ stand for the contribution of each branch to the global modulus. The overall viscosity *η* can be defined as Equation 3. The time-constant values were regularly distributed between the reciprocals of the highest (62.8 rad/s) and the lowest (0.628 rad/s) angular frequencies of the experimental dynamic modulus. The chosen number of modes was sufficiently high to obtain accurate fitting, but not too large to avoid inconsistent results (*e*.*g*., negative values of *β*_*α*_). Practically speaking, this led to three-time constants (m = 2), regularly spaced on a logarithmic scale between 5 × 10^−2^ s and 5 × 10^−1^ s.

Identification was achieved by solving the following minimization problem described by equation (3):

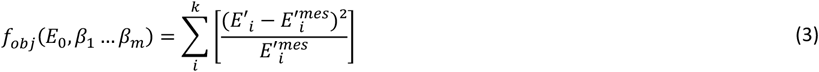

where 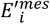 is the storage modulus obtained from the measured data and *E*′_*i*_ is the one computed with viscoelastic parameters. *k* is the number of measurements acquired during the frequency sweep compression test. The optimization procedure was performed by using the Microsoft Excel Solver (version 2016) with the Generalized Reduced Gradient (GRG) nonlinear solving method.

***Zymography assay***

## RESULTS AND DISCUSSION

The present study aims to describe the biological events which support the evolution of a 3D bioprinted construct from biomaterial seeded with fibroblasts up to constructs presenting progressive dermis development phases with typical physiological characteristics. These phases, identified as “maturation phases”, imply dynamic and concomitant biological mechanisms including fibroblasts proliferation, cell secretion and remodeling of extracellular matrix (ECM), cell death, and tissue contraction.

Here, the first step of our study consists of characterization and understanding of each “maturation phase” of a 3D bioprinted dermis construct. Then, an optimization and maturation-driven approach was evaluated thanks to the screening of construct architecture and composition as well as the addition of media supplements. In this case, the focus was to identify the biological mechanisms or parameters which could enhance tissue functionalities by controlling the extracellular matrix secretion and cell proliferation.

### 1 Description of bioprinted dermis development phases

A descriptive approach based on both destructive and non-destructive analyses was applied to monitor in depth the bioprinted constructs maturation phases for 34 days. Cellular proliferation was evaluated thanks both to cell counting after dissociation and live cell staining using calcein-green. Metabolic activity was evaluated through the monitoring of lactic acid release in the culture supernatant. Dermis construct matrix composition was evaluated thanks to histological analysis and collagen staining.

To evaluate the impact of cell proliferation on the bioprinted dermis evolution, an additional tissue shrinkage factor was evaluated by image analysis on ImageJ. The shrinkage factor (%) of the constructs was monitored over the different cultivation phases and plotted towards the cell density inside the construct evaluated thanks to destructive cell counting. Successive proliferation and maturation phases were described and will be listed below.

For the first fifteen days of cultivation (***Growth phase*, Figure 1-A**), cells proliferate within the constructs at a growth rate of 0.20 days^-1^ (µ_3D_), corresponding to a doubling time of 3.5 days. Such a proliferation rate is twice lower than the population doubling time (PDT_3D_) observed for fibroblasts cultivated in standard 2D flasks (µ_2D_: 0.40 days^-1^; PDT_2D_: 1.7 days). It is characterized by a progressive change in cell morphology, from spherical shape (day 5) to elongated shapes (> day 10). The metabolic activity (**Figure 1-A**) is also quite low compared to standard 2D cell metabolic activity as the expected specific lactic acid production yield Y_lact/X_ in such proliferation phase is of 1.9 g/(L.10^6^ cell).

Then, between 15^th^ and 19^th^ day of cultivation (***Death phase***), the 3D bioprinted dermis exhibit a second maturation phase. It is characterized by a stabilization of the growth rate for about 5 days before the onset of a 10-fold decrease of the cell number detected in the bioprinted constructs (**Figure 1-A**). On the metabolic behavior, no clear change is observed before the last maturation phase after the 20^th^ day of cultivation (***ECM Secretion***) where the cell metabolic activity is increased by 17 times, with specific lactic acid production rate reaching 33g/(L.10^6^cell) (**Figure 1-A**).

Interestingly, plotting total cell number as a function of cell density within the biomaterial allowed to identify these two breakpoints in the maturation phase. Indeed, on **Figure 1-B**, it is visible that after 15 days of growth, cells start to die with a decrease of the total cell number. Nevertheless, while cell number is reducing, on the contrary, the number of cells / g of biomaterial is increasing by 50%, then reaching 2.15 × 10^6^ cells/g, suggesting a reduction of the tissue construct volume over time. This allows reaching approximately 2200 cell/mm^3^ of constructs which falls within the 2100-4000 cell/mm^3^ described by Miller at al. [29] for dermis biopsies (**Figure 1-B**).

This was confirmed thanks to the analysis of hydrogel contraction. Indeed, macroscopic image analysis based on hydrogel contraction assays developed by Bell et al. (1979) allows to quantify such tissue shrinking [30]. Shrinkage factors were evaluated over the different maturation periods. It was demonstrated to reach 62% size reduction after 15 days of culture and over 79% at the end of the cultivation. This shrinkage results from two concomitant behaviors, first the fibroblast’s migration within the hydrogel which applies traction on the collagen fibers, second, the differentiation of fibroblasts into myofibroblasts. These cells are identified by α-SMA protein expression in their cytoskeleton, present a high level of collagen synthesis, and have a contractile function [33]. The presence of such cells with the apparition of collagen I synthesis was observed since day 15 in the dermis constructs (**Figure 1-C**). Such changes are coordinated with modifications of the bioprinted construct mechanical properties (**Figure 1-A**). Indeed, Young’s modulus (E0) remained stable at about 5.5 kPa during the ***Growth phase***. Then it is suddenly dropping to 2kPa during ***Death phase*** before increasing again during the ***ECM Secretion phase*** (from 19 and 34 days). Interestingly, it is also during this last phase that the bioprint construct viscosity is strongly increased from 0.2 to 3.8 kPa.s.

Microscopic observation and histological cuts allowed to confirm the successive events even more clearly along the three maturation phases identified. As described in **Figure 1-B**, viable cells observed thanks to calcein staining are homogeneously spread within the biomaterial for the first 15 days of cultivation. Cells are isolated, easily distinguished with a round shape up to day 5. Then, they are starting to spread, colonizing the whole bioprinted biomaterial. It is only from day 15 that fibroblasts density is visually increasing at the surface of the biomaterial. This is confirmed with histological analysis demonstrating densification of cells at the surface of the bioprinted dermis at the last cultivation stages **(Figure 1-B)**.

Such biological processes and phases have already been described in detail *in-vivo* and *in-vitro* during wound healing [31,32]. Thus, literature reports fibroblastic cells increase with the subsequent apparition of a new phenotype, i.e. myofibroblasts, characterized by specific cytoskeleton elements and the expression of α-SMA proteins. This cell population contributes to the production of granulation tissue and to wound contraction [33,34].

### 2 Bioprinted architecture and bioink composition impacting dermis development phases

Classical 3D bioprinted dermis present thickness between 0.4 and 2 mm [1,35,36] and mechanical strength of few hundred Pa [37]. Construct thickness used in literature has been driven by physiological considerations (mean thickness of the native dermis) but also by *in vitro* constraints such as nutriment availability within non-vascularized tissues [38]. Indeed, nutrient supply to cells was described to have a strong impact on tissue maturation and functionalization [39]. Similarly, biomaterial composition was demonstrated to have an impact on fibroblast proliferation and behavior [40]. In our case, two formulations of bioinks allowing good construct printability were evaluated (**formulation A-printed at 25°C**: 5% (w/v) of gelatin-2% (w/v) of Alginate-2% (w/v) of fibrinogen; **formulation B-printed at 28°C**: 10% (w/v) of gelatin-1% (w/v) of Alginate-2% (w/v) of fibrinogen). Such ratios modify the biomaterial mechanical properties, its internal porosity, and its degradability by cell metalloproteinases MMPs, while maintaining the bioink thinning behavior, highly advantageous for cell printing and protection from shear stress [41]. (see **Supplementary Figure 1)**

The impact on cell proliferation of the 3D bioprinted constructs architectures (thickness and porosity) and the biomaterial composition were investigated. **Figure 2-B** presents the two types of 3D architectures tested for the present study, i.e. bioprinted constructs of assembled bioink filaments of 400 µm diameter deposited either in a dense structure with no porosity (thick and thin constructs) or 300 µm gaps between filaments to generate porous architecture (porous constructs). Construct thickness was also investigated with 4 mm (thick) and 2 mm (thin) constructs. **Figure 2-A and 2-B** presents the growth profiles obtained for the different architectures and the two bioink formulations.

**Figure 2.**
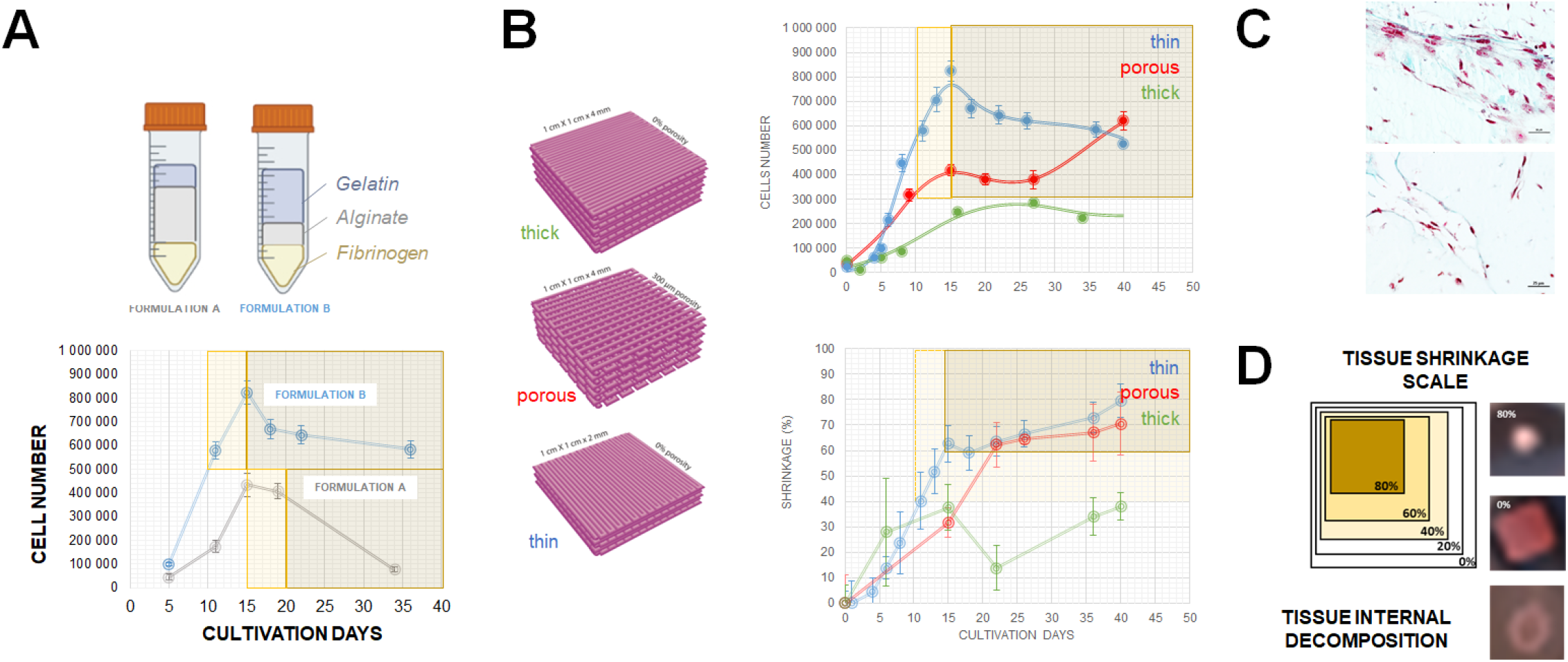
A- Impact of the biomaterial formulation on the growth profile with a thin 2mm architechture: formulation A (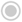) & B (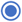). B- Growth profiles of bioprinted constructs depending on their architectures (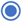 thin; 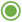 thick; 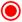 porous construct) prepared in formulation B. C-Histological staining demonstrating cell growth within the construct (D- Representation of the evaluation of construct size reduction (Shrinkage %). Evolution of the construct size reduction over the cultivation period.

#### Biomatrix composition

Similar growth trends were observed for both bioink formulations when bioprinted as a thin (2 mm) architecture configuration (**Figure 2-A)**. The three different phases (***growth, death***, and ***ECM secretion***) previously described were thereby conserved. Nevertheless, the growth capacity seems to be strongly increased in **formulation B**. Indeed, the presence of an increased amount of gelatin (10% over 5%) together with a reduced amount of alginate (1% versus 2%) allows to double the cell number in the bioprinted dermis construct after 20 days of culture. This also allows to reach 9 × 10^6^ cell/g which is more than 4 times the cell density achieved in formulation with less gelatin (Formulation A). Then, the growth rate of 0.51 days^-1^ and the doubling time of 1.4 days, demonstrate it was possible to recover the fibroblast growth capacities observed in static 2D cell culture. Microstructure’s description of the bioprinted materials demonstrates a differential microporosity observed on negative staining electron microscopy (see **Supplementary Figure 2**), correlated with differential biomechanical properties (**formulation A**-E0: 12.7kPa; **formulation B**-E0: 8.4kPa). Bioink composition allowed in our case to modulate the biomaterial microporosity and to recover optimal pore sizes like the one previously observed within collagen sponge for dermal constructs (50 to 120µm) [41].

#### Bioprinted architecture

Modifying the bioprinted construct architecture, while selecting the best bioink formulation (**formulation B**), allows distinguishing two maturation behaviors and the limiting factors impeding fibroblast proliferation and ECM secretion. The first set (thin and porous structures) of printed architectures exhibits active growth profiles and a maturation phase with tissue shrinkage and densification. In contrast, the second type of behavior observed for thick construct demonstrates reduced growth and biomaterial dissolution. Here the cell growth and activity is not sufficient to perform efficient remodeling of the biomaterial thus ending in complete destabilization of the construct.

Both thin and porous tissues demonstrate an active growth profile over the first 15 days (**Figure 2-B**) of cultivation with doubling time ranging from 1.4 days (thin) and 3.5 days (porous). After 40 days of culture, the shrinkage factor achieved for both constructs is similar, with 79% and 70% shrinkage factors for thin and porous tissues, respectively (see **Figure 2-B)**.

Both reached a cell density between 2-3 × 10^6^ cells/g when starts the shrinkage phase. In the case of thin tissue, this ***ECM secretion phase*** starts earlier than **formulation A** at 11 days of culture, thus indicating that cell density inside the bioprinted construct is critical for maturation onset.

On the contrary, thick constructs exhibit a population doubling time of 5.5 days with maximal cell density, which stabilizes after 15 days of cultivation at 5.6 × 10^5^ cells/g, which is 5 times lower than other constructs. In this case, the last maturation phase commonly associated with tissue shrinkage does not occur (construct shrinkage factor below 40%). The thick constructs started to soften until reaching complete destabilization after 20 days of cultivation.

A clear secretion of extracellular matrix was identified on the histological section colored with Masson’s Trichrome for both thin and porous architectures, as demonstrated in **Figure 2-C** and **Supplementary Figure 3**. The histological cuts also revealed the internal dermal construct cellularization (See **Supplementary Figure 3**). This later observation associated with the clear effect of construct geometry and porosity on fibroblasts confirms that media diffusion within the seeded biomaterial has a strong impact on cell proliferation.

Nutrient and oxygen transfer limitations were already described to occur as soon as the thickness of the biomaterial separating cells from the nutritive medium exceeds 300 µm [38]. In our case, thin and porous geometry allowed to have a maximum distance between cells and nutritive medium of respectively 1 mm and 200 µm. In the present study, growth limitations occur, only for 4 mm thick geometries where cells are drastically distanced from the culture medium.

### 3 Impact of cell growth factor and medium additives

Fibroblast proliferation and migration have been extensively studied and reviewed in the last four decades. It is largely accepted that multiple growth factors are secreted by fibroblastic cells and interplay with their proliferation and extracellular matrix secretion. Among them, within the Fibroblast Growth Factor (FGF) family, FGF-2 (or basic FGF), FGF-7, FGF-10, and FGF-21 are the ones especially impacting dermis migration and proliferation [25,42,43]. Epidermal growth factors (EGF) were also referenced to induce fibroblast proliferation.

To evaluate the impact of fibroblast secreted growth factors on their 3D proliferation and behavior, we performed a preliminary study in cultivating dermis constructs with a nutritive medium supplemented with 2D fibroblast cultures conditioned medium. Conditioned media correspond to 2D fibroblast culture medium collected at confluency after 7 days of growth. Nutritive DMEM medium was thus supplemented with 35% of conditioned medium and either used as a feeding medium or as part of the bioink formulation (**Figure 3-A**). The goal was to identify the impact of such growth factors on the proliferation rate. To evaluate the strongest effects, the **formulation A** providing the lowest growth in our previous studies was selected. This condition was chosen with a thin architecture of 2 mm. **Figure 3-A** presents the growth kinetics thanks to microscopic observations of calcein-stained viable cells at several time points.

**Figure 3.**
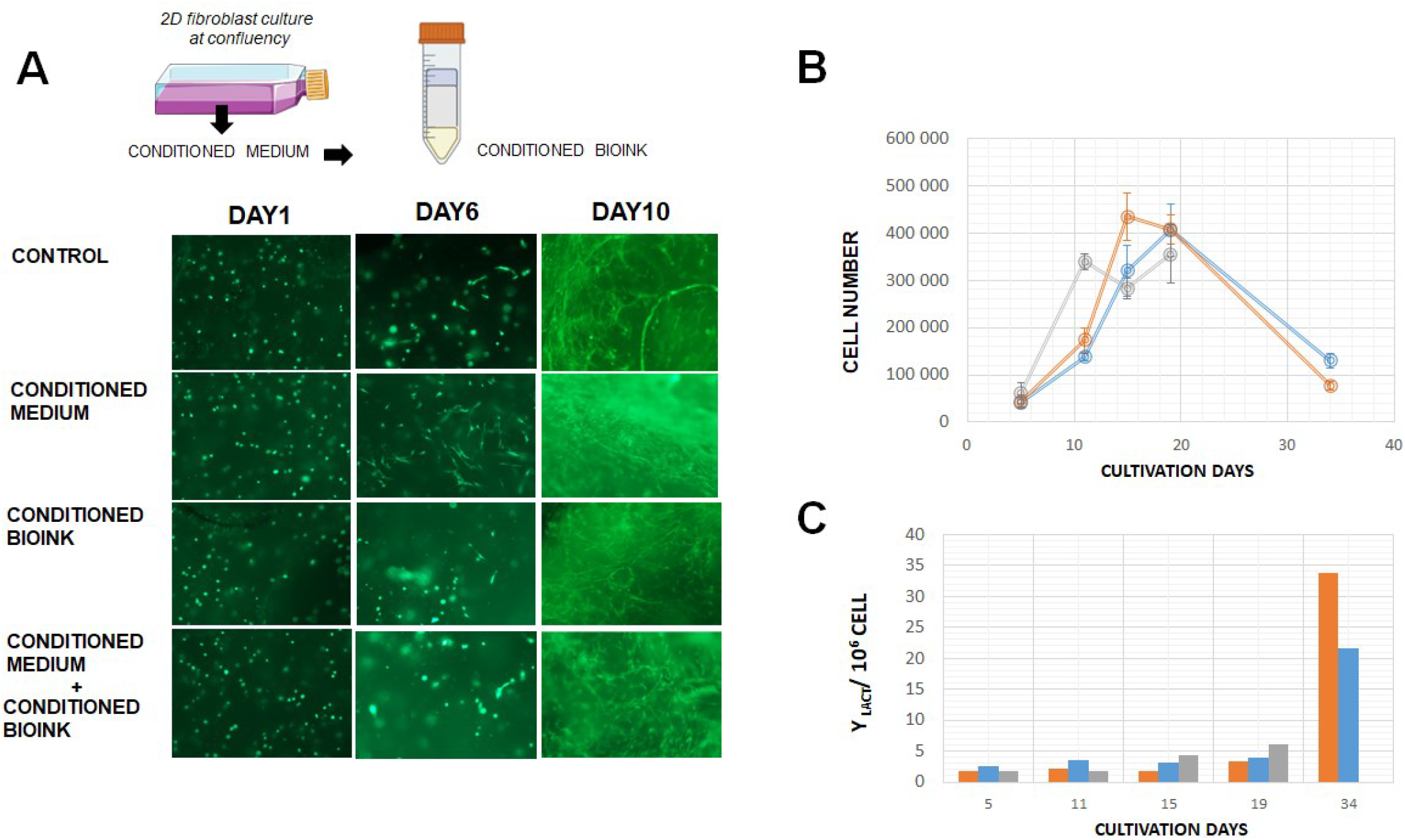
Impact of growth factors on bioprinted dermis’ fibroblast proliferation (thin construct 2mm). A- Impact of fibroblast conditioned medium on cell’s proliferation pattern (viable cell staining with calcein green). B- Growth profile of dermis fibroblast cells and C- lactate production rate in control condition (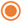) ; medium supplemented with 10ng/ml hEGF (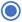) or 10ng/ml FGF-2 (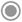).

Trends observed in **Figure 3-A** demonstrate that a conditioned medium strongly impacts cell proliferation during the first cultivation days when applied in the feeding medium. Indeed, as soon as the 6^th^ day of cultivation, fibroblasts present spread morphology within the biomatrix, while for the other cultivation conditions, this behavior occurs after 10 days of culture.

To go further in the comprehension of factors promoting 3D fibroblasts proliferation, two selected growth factors were supplemented to the cultivation media, human EGF (hEGF) and FGF-2. Molecule concentrations already reported in the literature [26] were chosen (10 ng/ml) with media renewal every 5 days. Growth profiles and lactate production rate are compared in **Figure 3-B** and **3-C**, respectively. As expected, FGF-2 seems to induce a fast fibroblast proliferation in 3D. The growth profile was twice faster than for other cultivation conditions (PDT_3D-FGF_ : 2.45 days). The counterpart of such rapid growth is the construct destabilization and its early dissolution after 19^th^ days of culture. On the contrary, the presence of hEGF does not strongly impact the cell growth profile. Similar cell count and metabolic behavior to the control condition were observed.

The evolution of the bioprinted construct mechanical properties are also good indicators to monitor the effect of these growth factor supplements. Here, in correlation with the fast-growing behavior of the fibroblasts in the presence of FGF2, a destabilization of the construct was easily identified by the drastic decrease of the tissue Young’s modulus after 11 days only (See **Supplementary Figure 4**). From a mechanical standpoint, the presence of hEGF has little effect on Young’s modulus until the ECM production phase. Then, its presence seems to hinder the achievement of high Young’s modulus (0.9kPa compared to 4.35kPa for control) and elasticity (Day 34). Thus, such observation implies that fastening the cell growth inside the tissues is insufficient and should be counteracted by cell-extracellular matrix secretion to preserve the tissue structure.

#### Extracellular Matrix Secretion and remodeling

As detailed above, in 3D culture conditions presenting fast growth rates, dermis tissue reconstruction holds on both cell growth and extracellular matrix production. Nevertheless, *in-vivo* fibroblast growth includes a first step consisting of extracellular matrix degradation thanks to the secretion, in their environment, of a family of Matrix Metalloproteinases (MMP) [17–19]. MMPs are a large family of enzymes able to degrade a wide range of biomolecules starting with ECM compounds [22], growth factors, growth factors inhibitors, but also membrane receptors, hormones, cytokines and other activators/inhibitors for several signaling pathways [44]. Skin dermis fibroblasts used for this study were characterized to secrete MMP-2 in 2D cultures (see **Supplementary Figure 5**). This enzyme is part of the gelatinase family [45]. Some recent studies showed that adding MMP inhibitors can reduce tissue contraction by avoiding fibroblasts’ migration and differentiation into myofibroblasts. They were also proved to reduce collagen I synthesis when fibroblasts are seeded onto collagen lattices [46].

*In vivo*, it was demonstrated that hepatic and pulmonary fibrosis is increased when high activity levels of MMP-2 and MMP-9 are detected whereas fibrosis was decreased after the administration of Batimastat® which inhibits MMP-2 and MMP-9 among other MMP [47,48]. Thus, the administration of Batimastat® decreased ECM secretion. Finally, Prinomastat® was described as an inhibitor of MMP 2, 3, 9, 13 and 14 [24] while Batimastat® only inhibits MMP 2, 3, and 9 but also MMP 1 and 8 [23].

We investigate the effect of conditions impacting extracellular matrix production. Here, vitamin C, known to stimulate collagen synthesis [49], and two pharmaceutical inhibitors of MMP, Batimastat®, and Prinomastat® were tested. Shrinkage, as well as cell growth and collagen secretion, were compared for each condition before (day 15) and after (day 22) the onset of the ECM secretion phase in **formulation B**. These experiments were also performed in 2mm thin construct. This is also the period where construct dissolution is occurring in the fast-growing behavior.

The macroscopic retractation due to media supplementation is not strongly different than the control for the different supplements. None of the conditions present a significative difference (p-value <0.05) with the control condition (**Figure 4-A**). A clear negative effect is visible on the growth rate when media are supplemented with MMP inhibitors. Indeed, Prinomastat® and Batimastat® demonstrate a total cell number in the construct reduced by half after 26 days of culture compared to the control condition and a growth rate close to zero (**Figure 4-B**). In conditions supplemented with MMP inhibitors, fibroblasts are not spreading through the construct internal zone (see Figure 4C). The hypothesis is here that no gelatin degradation occurred when MMPs were inhibited, leaving less porosity for the fibroblasts to develop. Thus, in both Prinomastat® and Batimastat® conditions, gelatin has been less degraded and ECM less neo-synthetized. This is consistent with literature which correlates the dermis fibroblast capacity to colonize a hydrogel and to replace it with their cellular matrix thanks to the secretion of MMP enzymes. It is also described in literature the role of collagen fragments released during fibroblasts degradation of ECM thanks to MMP-2 and MMP-9 on their proliferation capacity [50].

**Figure 4.**
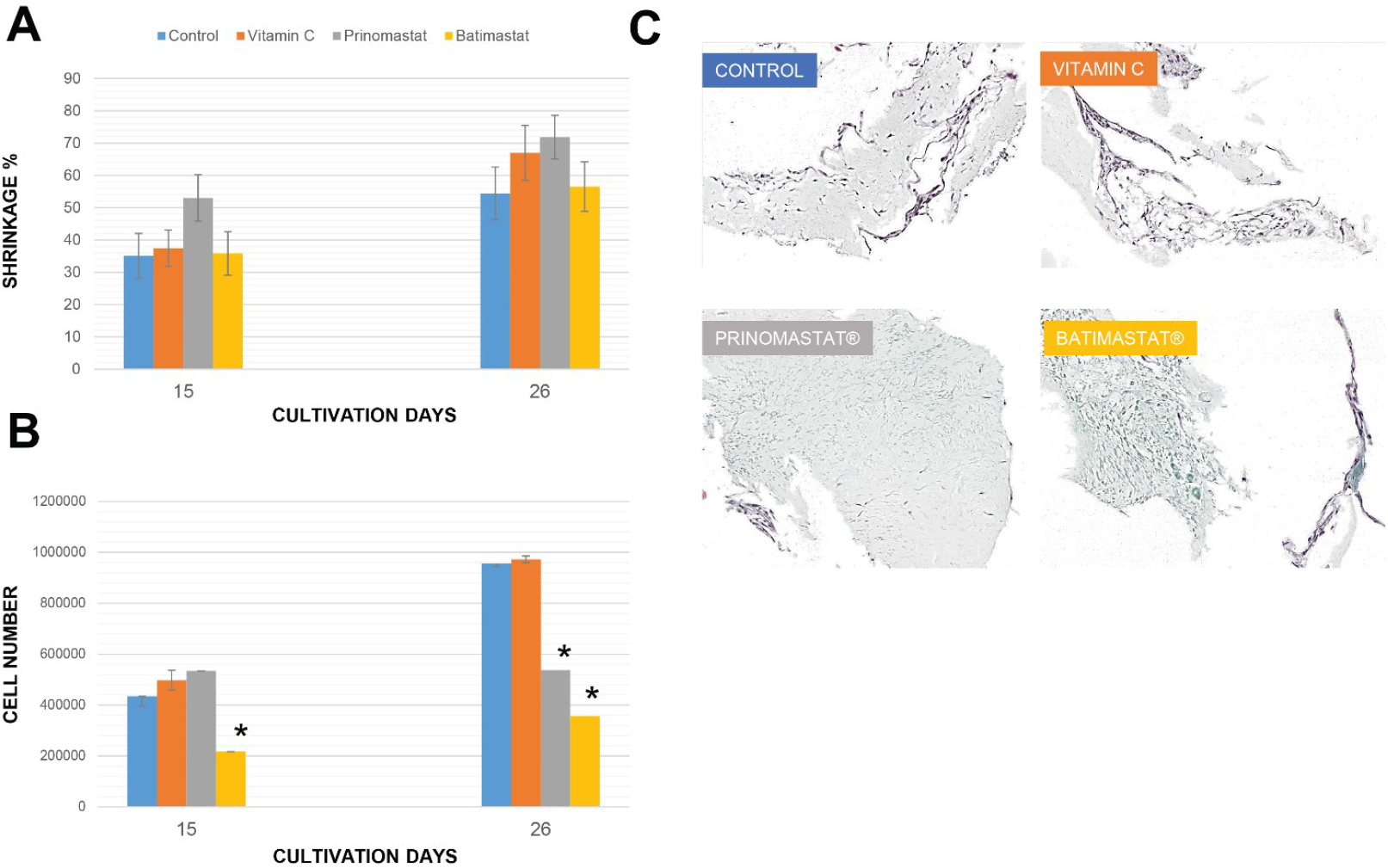
Evaluation of culture conditions on cell proliferation ECM secretion on thin 2mm constructs. A- Tissue shrinkage (%); B- Cell proliferation; C- Histological staining of ECM with Masson’s Trichrome (cells are colored in violet and collagen in green). Culture condition presented are control dermis construct cultivated without supplementation (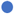), supplemented with Vitamin C (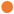), Prinomastat® (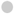), Batimastat® (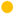). These TIMP and Vitamin C were added to the cultures every 2 days, and the culture media were completely renewed once a week in each culture condition with or without supplementation. Constructs were produced with Formulation B.

Finally, in the presence of vitamin C, cell growth was found to be like control condition and characterized by the colonization of the internal structure of the bioprint. Vitamin Cis described in literature as the co-factor of prolyl and lysyl hydroxylases taking part in collagen biosynthesis [51]. Nevertheless, in the present conditions, collagen synthesis enhancement was not confirmed thanks to histological observation (**Figure 4-C**). Previous studies described an increase in collagen I synthesis in ligament fibroblasts cultured with vitamin C only in presence of IGF-II [52]. Thus, vitamin C supplementation should be associated with IGF-II to further improve the dermis ECM biosynthesis.

## Conclusion

The present study is the first to describe in-depth the sequence and physiological behavior occurring during tissue development issued from a bioprinting strategy. We aimed to deeply detail and understand the parameters impacting cellular behavior and interaction with their 3D and nutritive environment to further allow for mastering and orientating such behavior. The strategy was applied to skin dermis constructs but could further be extrapolated to other bioprinted tissue. Thus, in this study, three development phases were identified that closely correspond to *in-vivo* wound healing steps: a ***Growth*** phase during which fibroblasts proliferate within the seeded biomaterial, a ***Death*** phase where fibroblasts differentiate into myofibroblasts and the total cell per construct decrease, an ***ECM secretion*** phase based on remodeling and matrix production. A tight balance must be found between providing appropriate conditions for cell growth and allowing for a concomitant extracellular matrix secretion to enable a bioprinted construct to reach the state of engineered dermis tissue. Indeed, we demonstrated that duration and intensity of these phases were strongly dependent upon the biomaterial properties (composition, architecture), the cell density, the cell feeding and the presence of ECM remodeling enhancers or inhibitors. This study also enabled settling a panel of destructive and non-destructive analytical methods which allow for the descriptions of behaviors of dermis cells and their environment.

## Figure Legends

**Supplementary Figure 1.**
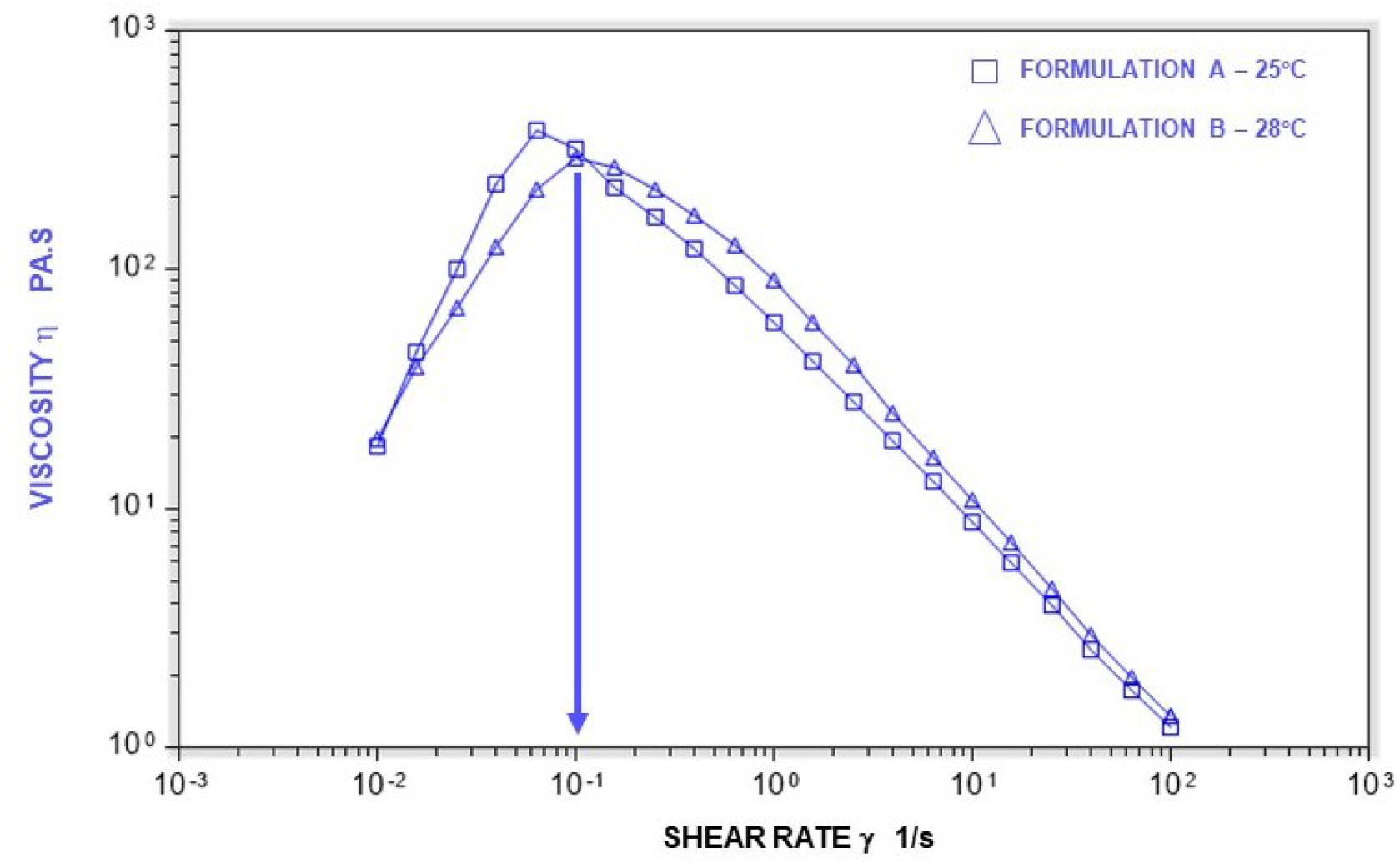
Rheological properties of bioink formulations A & B at their printing temperature. The arrow indicates the shear thinning behavior.

**Supplementary Figure 2.**
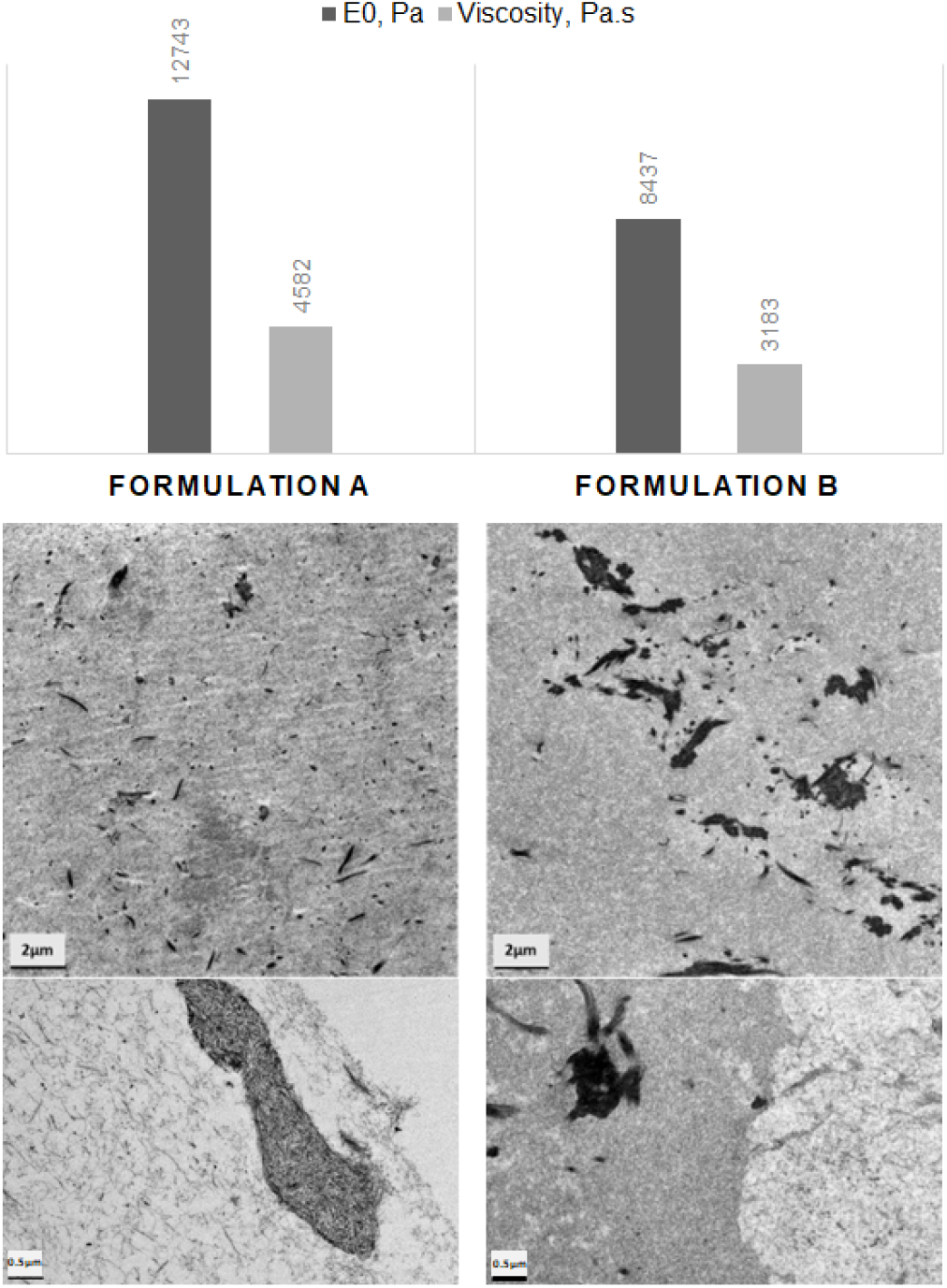
Characterization of mechanical properties (Young modulus and viscosity) and microstructures (Electron microscopy micrographs) of a bioprinted acellular construct.

**Supplementary Figure 3.**
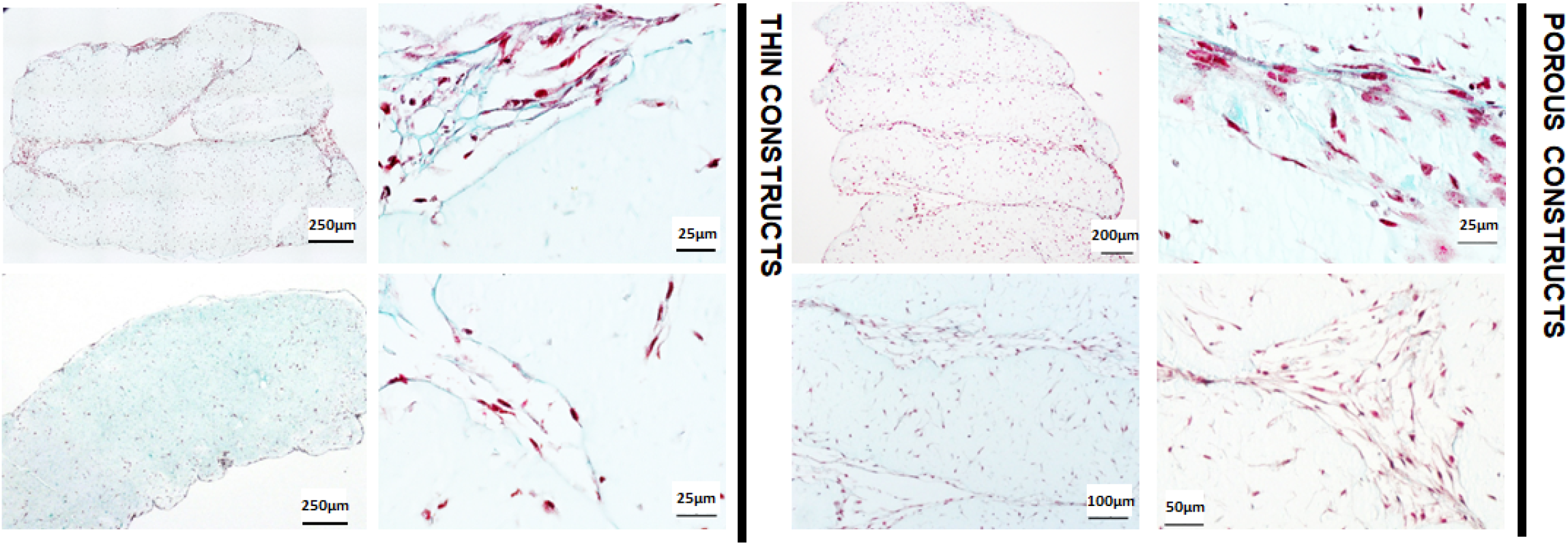
Histological analysis of cell repartition and collagen staining with Masson Trichrome. (Pink: cell nucleus; Blue-Green : collagen) for porous and thin architectures.

**Supplementary Figure 4.**
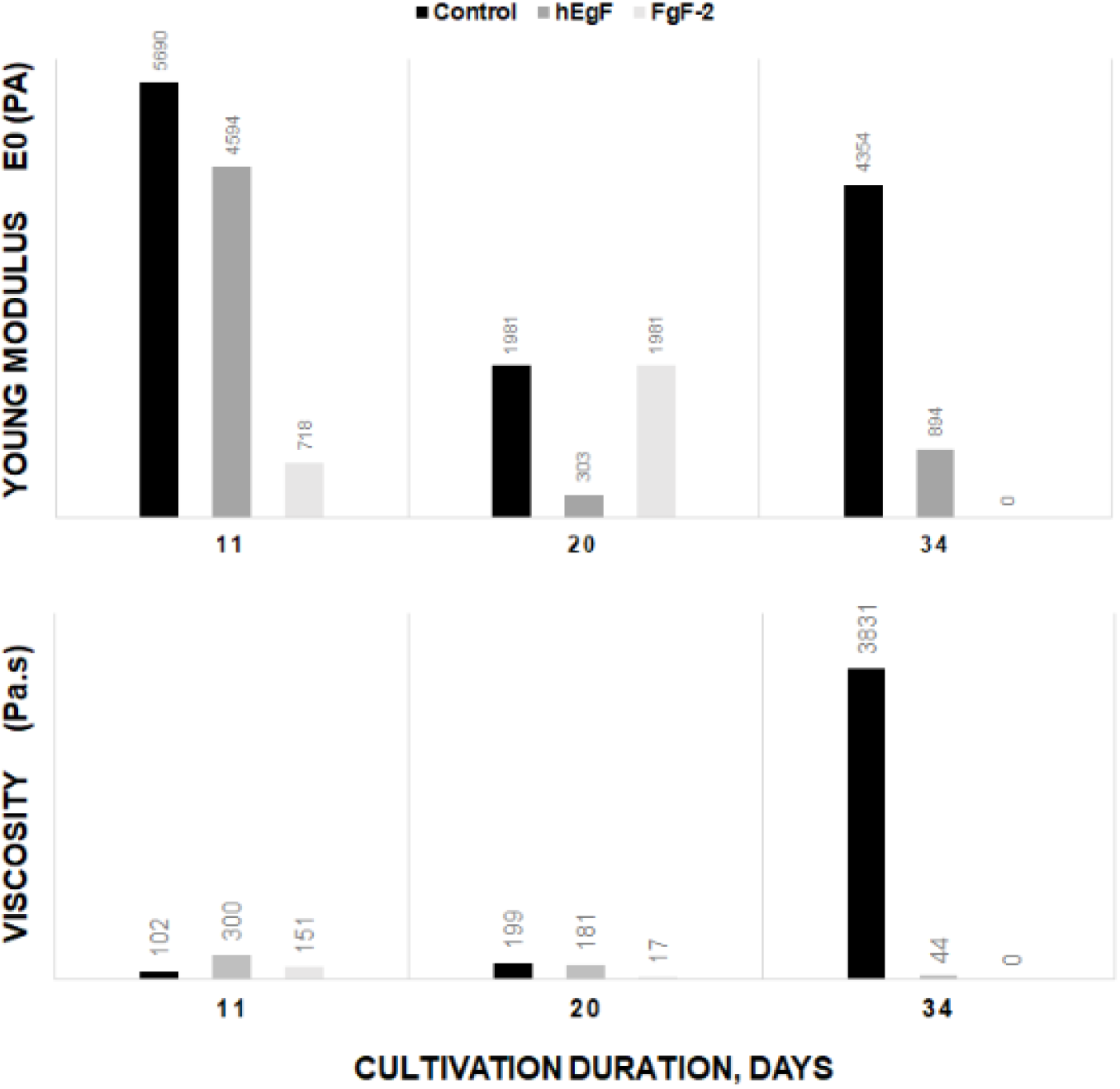
Mechanical properties of bioprinted constructs: Control (black), hEGF (Gray), FGF-2 (light grey). All conditions were evaluated on bioink formulation A.

**Supplementary Figure 5.**
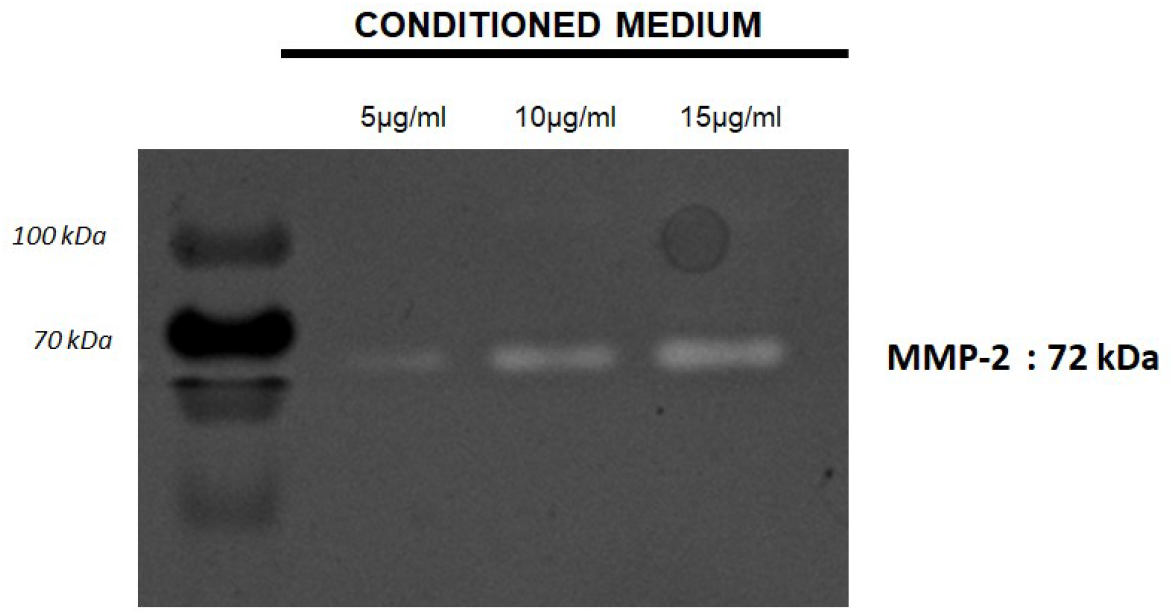
Zymography of fibroblasts conditioned medium collected after 7 days culture in adherence to reach confluency. Protein content of conditioned medium was quantified by BCA assay to load respectively 5, 10 or 15µg/ml per well. MMP-2 enzyme was the sole MMP identified in conditioned medium.

## Literature

[1] L.J. Pourchet, A. Thepot, M. Albouy, E.J. Courtial, A. Boher, L.J. Blum, C.A. Marquette, Human Skin 3D Bioprinting Using Scaffold-Free Approach, Adv. Healthc. Mater. (2016) 1601101. https://doi.org/10.1002/adhm.201601101.

[2] F. Salaris, A. Rosa, Construction of 3D in vitro models by bioprinting human pluripotent stem cells: Challenges and opportunities, Brain Res. 1723 (2019) 146393. https://doi.org/10.1016/j.brainres.2019.146393.

[3] E. Masaeli, V. Forster, S. Picaud, F. Karamali, M.H. Nasr-Esfahani, C. Marquette, Tissue engineering of retina through high resolution 3-dimentional inkjet bioprinting, Biofabrication. (2020) 1–32.

[4] S. V. Murphy, A. Atala, 3D bioprinting of tissues and organs, Nat. Biotechnol. 32 (2014) 773–785. https://doi.org/10.1038/nbt.2958.

[5] H. Huang, H. Xu, J. Zhang, Current Tissue Engineering Approaches for Cartilage Regeneration, in: D. D. Nikolopoulos, G. K. Safos, K. Dimitrios (Eds.), Cartil. Tissue Eng. Regen. Tech., IntechOpen, 2019.

[6] I.C.P. Rodrigues, A. Kaasi, R. Maciel Filho, A.L. Jardini, L.P. Gabriel, Cardiac tissue engineering: current state-of-the-art materials, cells and tissue formation, Einstein (São Paulo). 16 (2018). https://doi.org/10.1590/s1679-45082018rb4538.

[7] G. Mazza, W. Al-Akkad, K. Rombouts, M. Pinzani, Liver tissue engineering: From implantable tissue to whole organ engineering, Hepatol. Commun. 2 (2018) 131–141. https://doi.org/10.1002/hep4.1136.

[8] G. Cesare, Muscle tissue engineering, in: Regen. Eng. Musculoskelet. Tissues Interfaces, Elsevier, 2015: pp. 239–268.

[9] C. Auxenfans, V. Menet, Z. Catherine, H. Shipkov, P. Lacroix, M. Bertin-Maghit, O. Damour, F. Braye, Cultured autologous keratinocytes in the treatment of large and deep burns: A retrospective study over 15 years, Burns. 41 (2015) 71–79. https://doi.org/10.1016/j.burns.2014.05.019.

[10] A.W. Feinberg, Engineered tissue grafts: opportunities and challenges in regenerative medicine: Engineered tissue grafts, Wiley Interdiscip. Rev. Syst. Biol. Med. 4 (2012) 207–220. https://doi.org/10.1002/wsbm.164.

[11] M. Roger, N. Fullard, L. Costello, S. Bradbury, E. Markiewicz, S. O’Reilly, N. Darling, P. Ritchie, A. Määttä, I. Karakesisoglou, G. Nelson, T. von Zglinicki, T. Dicolandrea, R. Isfort, C. Bascom, S. Przyborski, Bioengineering the microanatomy of human skin, J. Anat. 234 (2019) 438–455. https://doi.org/10.1111/joa.12942.

[12] N. Sigaux, L. Pourchet, M. Albouy, A. Thépot, C. Marquette, Is 3D Bioprinting the Future of Reconstructive Surgery?, Plast. Reconstr. Surg. - Glob. Open. 5 (2017) e1246. https://doi.org/10.1097/GOX.0000000000001246.

[13] T.M. Brown, K. Krishnamurthy, Histology, Dermis, in: StatPearls, StatPearls Publishing, Treasure Island (FL), 2020.

[14] O. A., Aging of the skin connective tissue: how to measure the biochemical and mechanical properties of aging dermis., Photodermatol Photoimmunol Photomed. 10 (1994) 47–52.

[15] M.A. Cole, T. Quan, J.J. Voorhees, G.J. Fisher, Extracellular matrix regulation of fibroblast function: redefining our perspective on skin aging, J. Cell Commun. Signal. 12 (2018) 35–43. https://doi.org/10.1007/s12079-018-0459-1.

[16] L. Pourchet, E. Petiot, C. Loubière, E. Olmos, M. Dos Santos, A. Thépot, B.J.B.J.B.J. Loïc, C.A.C.A. Marquette, Large 3D bioprinted tissue: Heterogeneous perfusion and vascularization, Bioprinting. 13 (2019) 1–7. https://doi.org/10.1016/j.bprint.2018.e00039.

[17] T. Tsuruda, L.C. Costello-Boerrigter, J. Burnett John C., Matrix Metalloproteinases: Pathways of Induction by Bioactive Molecules, Heart Fail. Rev. 9 (2004) 53–61. https://doi.org/10.1023/B:HREV.0000011394.34355.bb.

[18] V. Pelmenschikov, P.E.M. Siegbahn, Catalytic Mechanism of Matrix Metalloproteinases: Two-Layered ONIOM Study, Inorg. Chem. 41 (2002) 5659–5666. https://doi.org/10.1021/ic0255656.

[19] D. Rodríguez, C.J. Morrison, C.M. Overall, Matrix metalloproteinases: What do they not do? New substrates and biological roles identified by murine models and proteomics, Biochim. Biophys. Acta - Mol. Cell Res. 1803 (2010) 39–54. https://doi.org/10.1016/j.bbamcr.2009.09.015.

[20] A. Desmoulière, C. Chaponnier, G. Gabbiani, Tissue repair, contraction, and the myofibroblast, Wound Repair Regen. 13 (2005) 7–12. https://doi.org/10.1111/j.1067-1927.2005.130102.x.

[21] T. Grzela, A. Krejner, M. Litwiniuk, Matrix metalloproteinases in the wound microenvironment: therapeutic perspectives, Chronic Wound Care Manag. Res. (2016) 29. https://doi.org/10.2147/CWCMR.S73819.

[22] H. Birkedal-Hansen, W.G.I. Moore, M.K. Bodden, L.J. Windsor, B. Birkedal-Hansen, A. DeCarlo, J.A. Engler, Matrix Metalloproteinases: A Review, Crit. Rev. Oral Biol. Med. 4 (1993) 197–250. https://doi.org/10.1177/10454411930040020401.

[23] U. Mirastschijski, R. Schnabel, J. Claes, W. Schneider, M.S. Ågren, C. Haaksma, J.J. Tomasek, Matrix metalloproteinase inhibition delays wound healing and blocks the latent transforming growth factor-β1-promoted myofibroblast formation and function, Wound Repair Regen. 18 (2010) 223–234. https://doi.org/10.1111/j.1524-475X.2010.00574.x.

[24] R. Scatena, Prinomastat, a hydroxamate-based matrix metalloproteinase inhibitor. A novel pharmacological approach for tissue remodelling-related diseases, Expert Opin. Investig. Drugs. 9 (2000) 2159–2165. https://doi.org/10.1517/13543784.9.9.2159.

[25] Y.H. Song, Y.T. Zhu, J. Ding, F.Y. Zhou, J.X. Xue, J.H. Jung, Z.J. Li, W.Y. Gao, Distribution of fibroblast growth factors and their roles in skin fibroblast cell migration, Mol. Med. Rep. 14 (2016) 3336– 3342. https://doi.org/10.3892/mmr.2016.5646.

[26] Y.Y. Jia, J.Y. Zhou, Y. Chang, F. An, X.W. Li, X.Y. Xu, X.L. Sun, C.Y. Xiong, J.L. Wang, Effect of optimized concentrations of basic fibroblast growth factor and epidermal growth factor on proliferation of fibroblasts and expression of collagen: Related to pelvic floor tissue regeneration, Chin. Med. J. (Engl). 131 (2018) 2089–2096. https://doi.org/10.4103/0366-6999.239301.

[27] J.T. Daniels, A.D. Cambrey, N.L. Occleston, Q. Garrett, R.W. Tarnuzzer, G.S. Schultz, P.T. Khaw, Matrix metalloproteinase inhibition modulates fibroblast-mediated matrix contraction and collagen production in vitro, Investig. Ophthalmol. Vis. Sci. 44 (2003) 1104–1110. https://doi.org/10.1167/iovs.02-0412.

[28] S. Pragnere, N. El Kholti, L. Gudimard, L. Essayan, C. Marquette, E. Petiot, C. Pailler-Mattei, Quantification of cell contractile behavior based on non-destructive macroscopic measurement of tension forces on bioprinted hydrogel, J. Mech. Behav. Biomed. Mater. 134 (2022) 105365. https://doi.org/https://doi.org/10.1016/j.jmbbm.2022.105365.

[29] C.C. Miller, G. Godeau, C. Lebreton-DeCoster, A. Desmouliere, B. Pellat, L. Dubertret, B. Coulomb, A. Desmoulière, B. Pellat, L. Dubertret, B. Coulomb, Validation of a morphometric method for evaluating fibroblast numbers in normal and pathologic tissues, Exp. Dermatol. 12 (2003) 403– 411. https://doi.org/10.1034/j.1600-0625.2003.00023.x.

[30] E. Bell, B. Ivarsson, C. Merrill, Production of a tissue-like structure by contraction of collagen lattices by human fibroblasts of different proliferative potential in vitro, Proc. Natl. Acad. Sci. U. S. A. 76 (1979) 1274–1278. https://doi.org/10.1073/pnas.76.3.1274.

[31] A.L. Rippa, E.P. Kalabusheva, E.A. Vorotelyak, Regeneration of dermis: Scarring and cells involved, Cells. 8 (2019). https://doi.org/10.3390/cells8060607.

[32] P. Rossio-Pasquier, D. Casanova, A. Jomard, M. Démarchez, Wound healing of human skin transplanted onto the nude mouse after a superficial excisional injury: human dermal reconstruction is achieved in several steps by two different fibroblast subpopulations, Arch. Dermatol. Res. 291 (1999) 591–599. https://doi.org/10.1007/s004030050460.

[33] A. Desmoulière, C. Chaponnier, G. Gabbiani, Perspective Article: Tissue repair, contraction, and the myofibroblast, Wound Repair Regen. 13 (2005) 1067–1927.

[34] I.A. Darby, T.D. Hewitson, Fibroblast Differentiation in Wound Healing and Fibrosis, 257 (n.d.) 143–179. https://doi.org/10.1016/S0074-7696(07)57004-X.

[35] P. Admane, A.C. Gupta, P. Jois, S. Roy, C. Chandrasekharan Lakshmanan, G. Kalsi, B. Bandyopadhyay, S. Ghosh, Direct 3D bioprinted full-thickness skin constructs recapitulate regulatory signaling pathways and physiology of human skin, Bioprinting. 15 (2019) e00051. https://doi.org/10.1016/j.bprint.2019.e00051.

[36] K. Derr, J. Zou, K. Luo, M.J. Song, G.S. Sittampalam, C. Zhou, S. Michael, M. Ferrer, P. Derr, Fully Three-Dimensional Bioprinted Skin Equivalent Constructs with Validated Morphology and Barrier Function, Tissue Eng. - Part C Methods. 25 (2019) 334–343. https://doi.org/10.1089/ten.tec.2018.0318.

[37] M. Dussoyer, E.J. Courtial, M. Albouy, A. Thépot, M. Dussoyer, E.J. Courtial, M. Albouy, A. Thépot, M. Dos Santos, M. Dussoyer, E.J. Courtial, M. Albouy, A. Thépot, M. Dos Santos, C.A. Marquette, Mechanical properties of 3D bioprinted dermis : characterization and improvement To cite this version : HAL Id : hal-03027584 Mechanical properties of 3D bioprinted dermis : characterization and improvement, (2020).

[38] Y.M. Khong, J. Zhang, S. Zhou, C. Cheung, K. Doberstein, V. Samper, H. Yu, Novel Intra-Tissue Perfusion System for Culturing Thick Liver Tissue, Tissue Eng. 13 (2007) 2345–2356. https://doi.org/10.1089/ten.2007.0040.

[39] B. Porter, R. Zauel, H. Stockman, R. Guldberg, D. Fyhrie, 3-D computational modeling of media flow through scaffolds in a perfusion bioreactor, J. Biomech. 38 (2005) 543–549. https://doi.org/10.1016/j.jbiomech.2004.04.011.

[40] K. Bott, Z. Upton, K. Schrobback, M. Ehrbar, J.A. Hubbell, M.P. Lutolf, S.C. Rizzi, The effect of matrix characteristics on fibroblast proliferation in 3D gels, Biomaterials. 31 (2010) 8454–8464. https://doi.org/10.1016/j.biomaterials.2010.07.046.

[41] S. Lili, B. François, D. Odile, C. Christian, Chracterization of skin reconstructed on a chitosan-cross-linked Collagen-glycosaminoglycan matrix, Ski. Pharmacol. 3 (1990) 107–114.

[42] D.M. Ornitz, N. Itoh, Protein family review: Fibroblast growth factors, Genome Biol. 2 (2001) reviews3005.1-3005.12. http://genomebiology.com/2001/2/3/reviews/3005.

[43] A. Yu, Y. Matsuda, A. Takeda, E. Uchinuma, Y. Kuroyanagi, Effect of EGF and bFGF on fibroblast proliferation and angiogenic cytokine production from cultured dermal substitutes, J. Biomater. Sci. Polym. Ed. 23 (2012) 1315–1324. https://doi.org/10.1163/092050611X580463.

[44] M.D. Sternlicht, Z. Werb, How Matrix Metalloproteinases Regulate Cell Behavior, Annu. Rev. Cell Dev. Biol. 17 (2001) 463–516. https://doi.org/10.1146/annurev.cellbio.17.1.463.

[45] M.K. Winkler, J.L. Fowlkes, Metalloproteinase and growth factor interactions: do they play a role in pulmonary fibrosis?, Am. J. Physiol. Cell. Mol. Physiol. 283 (2002) L1–L11. https://doi.org/10.1152/ajplung.00489.2001.

[46] J.T. Daniels, A.D. Cambrey, N.L. Occleston, Q. Garrett, R.W. Tarnuzzer, G.S. Schultz, P.T. Khaw, Metalloproteinase Inhibition Modulates Fibroblast-Mediated Matrix Contraction and Collagen Production In Vitro, Investig. Opthalmology Vis. Sci. 44 (2003). https://doi.org/10.1167/iovs.02-0412.

[47] M. Corbel, S. Caulet-Maugendre, N. Germain, S. Molet, V. Lagente, E. Boichot, Inhibition of bleomycin-induced pulmonary fibrosis in mice by the matrix metalloproteinase inhibitor batimastat, J. Pathol. 193 (2001) 538–545. https://doi.org/10.1002/path.826.

[48] B. Vaillant, M.G. Chiaramonte, A.W. Cheever, P.D. Soloway, T.A. Wynn, Regulation of Hepatic Fibrosis and Extracellular Matrix Genes by the Th Response: New Insight into the Role of Tissue Inhibitors of Matrix Metalloproteinases, J. Immunol. 167 (2001) 7017–7026. https://doi.org/10.4049/jimmunol.167.12.7017.

[49] S. Murad, D. Grove, K.A. Lindberg, G. Reynolds, A. Sivarajah, S.R. Pinnell, Regulation of collagen synthesis by ascorbic acid., Proc. Natl. Acad. Sci. 78 (1981) 2879–2882. https://doi.org/10.1073/pnas.78.5.2879.

[50] A. Kisling, R.M. Lust, L.C. Katwa, What is the role of peptide fragments of collagen I and IV in health and disease?, Life Sci. 228 (2019) 30–34. https://doi.org/https://doi.org/10.1016/j.lfs.2019.04.042.

[51] T. Duarte, M. Cooke, G. Jones, Gene expression profiling reveals new protective roles for vitamin C in human skin cells, Free Radic. Biol. Med. 46 (2009) 78–87. https://doi.org/10.1016/j.freeradbiomed.2008.09.028.

[52] J.E. Moreau, J. Chen, D.S. Bramono, V. Volloch, H. Chernoff, G. Vunjak-Novakovic, J.C. Richmond, D.L. Kaplan, G.H. Altman, Growth factor induced fibroblast differentiation from human bone marrow stromal cells in vitro, J. Orthop. Res. 23 (2005) 164–174. https://doi.org/10.1016/j.orthres.2004.05.004.

